# Transposon-derived transcription factors across metazoans

**DOI:** 10.1101/2022.12.18.520930

**Authors:** Krishanu Mukherjee, Leonid L. Moroz

## Abstract

Transposable elements (TE) could serve as sources of new transcription factors (TFs) in plants and some other model species, but such evidence is lacking for most animal lineages. Here, we discovered multiple independent co-options of TEs to generate 788 TFs across Metazoa, including all early-branching animal lineages. Six out of ten super-families of known TEs (ZBED, CENPB, FHY3, HTH-Psq, THAP, and FLYWCH) were recruited as representatives of nine phyla. The most extensive convergent domestication of TE-derived TFs occurred in the hydroid polyps, polychaete worms, cephalopods, oysters, and sea slugs. Phylogenetic reconstructions showed species-specific clustering and lineage-specific expansion; none of the identified TE-derived TFs revealed homologs in their closest neighbors. Together, our study established a framework for categorizing TE-derived TFs and informing the origins of novel genes across phyla.

## Introduction

Transposable elements (TEs) or transposons, identified by Barbara McClintock in the 1940-the 50s, are now recognized as regulatory elements ^1^, which control ∼25% of human genes ^2^ (e.g., via promoters). TEs are also major constituents of all eukaryotic genomes, frequently occupying from 20% to more than 70% of genomes. The inherent ability of TEs to self-replicate, move and mutate transformed the initial assessment of TEs as ‘selfish gene’ parasites and ‘junk DNA’ into powerful evolutionary forces ^3^. The process of suppressing parasitic self-propagation properties is called molecular domestication or exaptation ^3-5^.

A domesticated TE-derived gene regulator can benefit the host and be an adaptive advantage ^1,3,4,6^. The TE-associated domestication events can be sources of novel genes ^3^, ncRNAs, microRNAs, etc. ^7-11^. There are multiple examples of such beneficial domestication events, and the scope of this process is expanding with sequenced genomes ^2-4,6,12,13^. There are also examples of convergent domestication, reflecting TE’s nature ^14,15^. For example, the emergence of the placenta from the TE-derived *Syncytin* gene in mammals and lizards occurred through two independent occurrences of TE domestication; it is portrayed as a classic example of convergent evolution ^3,16,17^.

Perhaps, the most critical domestication episodes associated with the rise of biological novelties are the recruitments of TEs in the evolution of transcription factors (TFs) known as master regulators of gene expression across Metazoa ^18,19^, including body patterning ^20,21^ and cell fate commitment ^22,23^. Mechanisms of the origins and lineage-specific TF gene expansion are primarily unknown. A classical hypothesis implies ancestral TF gene duplication, followed by the divergence of the duplicated gene ^24^. However, this scenario does not apply to the TFs that are solely organism-specific and have no *bona fide* one-to-one orthologs in closest relatives. The complementary scenario is the origin of TFs with the contribution of TEs. DNA-binding properties of TEs, in particular the evidence that TEs contain TF-binding sites, perfectly match structural genome constraints as a potential ‘pre-adaptation’ and sources to form novel cis-regulatory elements and TFs. Thus, the incorporation of non-coding and new TF genes into existing transcriptional networks ^25^ can also lead to the origins of new functions and transformative biological innovations, as well as the diversification of both genes and forms.

The most notable examples of TE-derived TFs came from plants ^11,26^ and such model animal species as insects (e.g., *Drosophila* ^3,15,27^) or vertebrates ^28-33^. In humans, there are examples of TE-derived TFs ^2,13,34^ and contributions of TE to diseases ^35^. However, the broad comparative scope of these events is less explored, with little knowledge about the majority of animal phyla.

Practically nothing is known about the most diverse bilaterian lineage– Lophotrochozoa. This clade consists of more than a dozen phyla ^36^, including Mollusca– the second most species-rich phylum and one of the most diverse groups of animals ^37^. The evidence of TE domestication events outside Bilateria in four other basal metazoan lineages (Ctenophora, Porifera, Placozoa, and Cnidaria) is also lacking.

Here, we generated a catalog of TE-derived TFs across Metazoa and proposed independent co-option of six out of ten superfamilies of TEs to create hundreds of TFs in all early-branching animal lineages.

## Results & Discussion

### 1. Mosaic distribution and parallel evolution of transposon-derived transcription factors across metazoans

We first curated a complete dataset of transcription factors (TFs) encoded in representatives of four animal phyla with the sequenced genomes, including two bilaterians (*Aplysia* and *Octopus*), one ctenophore (*Pleurobrachia*), a sponge (*Amphimedon*), and a placozoan (*Trichoplax*). We used the most completed and annotated dataset of 1600 TFs encoded in the human genome to represent the deuterostomes clade ^38^ and 755 predicted sequence-specific TFs in *Drosophila*, the model representative of the Ecdysozoa clade, as the initial queries ^39^. Utilizing these datasets, we identified that the sea slug *Aplysia* genome encodes 824 transcription factors. Similarly, using all *Aplysia, Drosophila*, and human TFs as queries, we identified the complete repertoire of TFs encoded in the *Octopus bimaculoides* and the other three (*Trichoplax, Amphimedon, Pleurobrachia*) basal metazoan genomes.

Next, we identified TF families in these four animal phyla that have undergone lineage-specific TFs gene expansions including the ones that have originated through tandem duplications. To our surprise, we found that the class II DNA transposable elements (TEs) were primarily associated with species-specific TFs family gene expansion (**Fig. 1)**. There are ten super-families of Class II TEs, and are known to use the ‘*cut-and-paste*’ mechanism for transposition from one position in the genome to another ^6,40^. Representatives of each of these subfamilies of proteins were used as a query to screen for TE-derived TFs across nine metazoan phyla (**Fig. 1, Table 1S, Table 2S)**. We determined that six of these TEs super-families could be independently recruited into the metazoan TFs: ZBED, CENPB, FHY3, HTH-Psq, THAP, and FLYWCH (**Fig. 1)**. Phylogenetic reconstruction suggested independent recruitment due to the absence of a ‘one-to-one’ homolog in the closest species (**Fig. 2)**. The domain organization of newly identified TE-derived metazoan TFs (summarized in **Fig. 3**), also revealed the presence of transposon-like components within the protein-coding open reading frames (ORFs). The occurence of TEs components within the TFs was further supported by sequence similarity searches against the *de novo* assembled transcriptome (RNA-Seq) dataset (https://neurobase.rc.ufl.edu).

**Table 1.** The total number of TE-derived TFs identified in this study.

| GENE/TFS | # | COMMENTS, THE FIRST DESCRIPTION | TOP 3-4 SPECIES WITH THE EXPANSION OF A GIVEN TF FAMILY |
| --- | --- | --- | --- |
| ZBED | 71 | - 1<sup>st</sup> for Lophotrochozoa | *Aplysia* (10), *Amphimedon* (13), ***Hydra*** (15) |
| CENPB | 121 | - 1<sup>st</sup> for Lophotrochozoa | ***Aplysia*** (14), *Homo* (12), *Octopus* (7) |
| FHY3 | 23 | - 1<sup>st</sup> for Metazoa | ***Aplysia*** (3), *Amphimedon* (3), *Lingula* (2), *Octopus* (1) |
| HTH-PSQ | 136 | - 1<sup>st</sup> for Lophotrochozoa | *Aplysia* (16), ***Hydra*** (43), *Octopus* (12) |
| THAP | 370 | - 1<sup>st</sup> for Lophotrochozoa | ***Capitella*** (87), *Hydra* (73), *Crassostrea* (58) |
| FLYWCH | 67 | - 1<sup>st</sup> for Ctenophora & Lophotrochozoa | Expansion in Ctenophores, |
| TOTAL | 788 |  |  |

**Figure 1:**
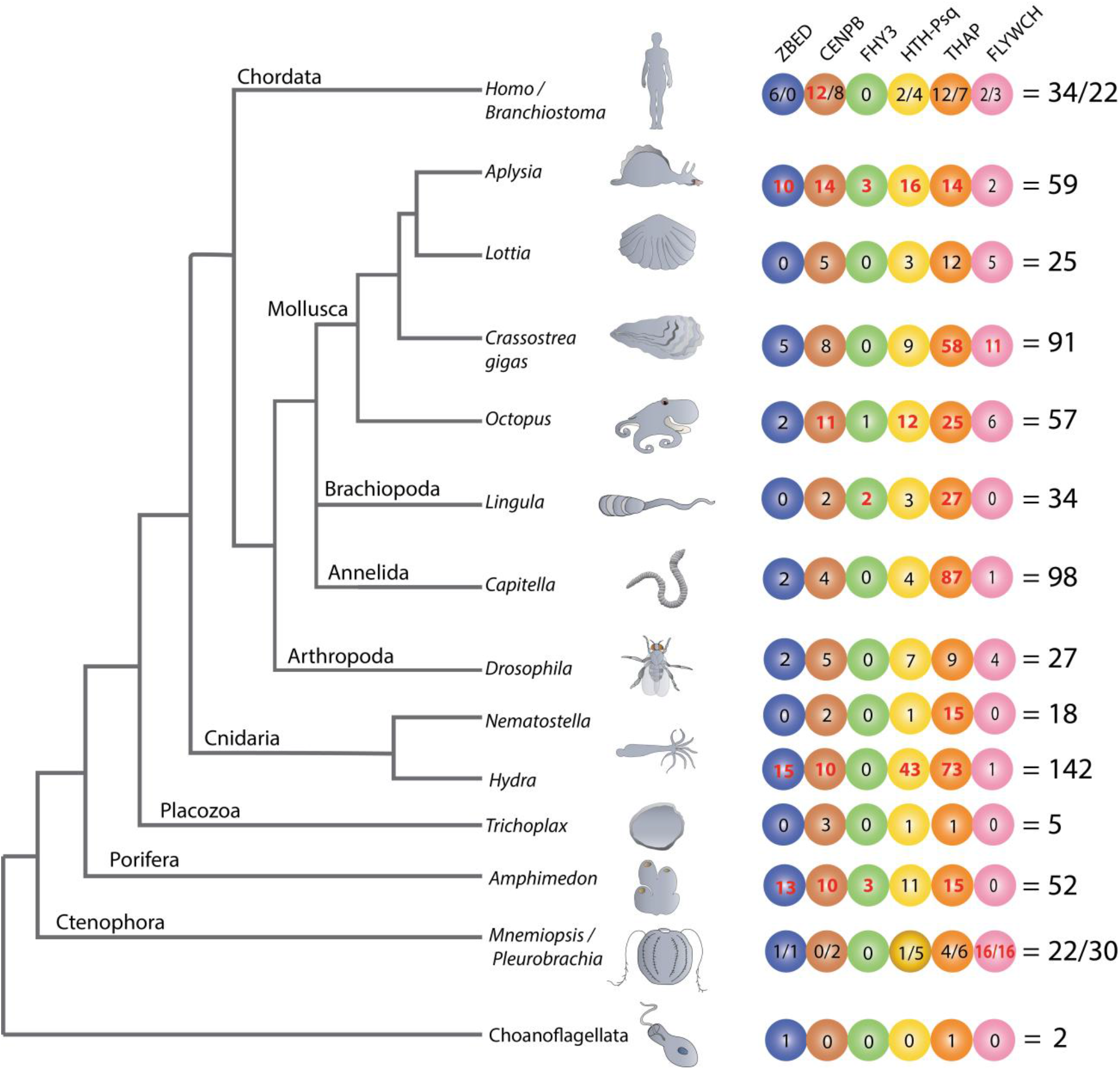
Transposon-derived transcription factors across metazoans. The diagram shows Lineage-specific expansion and mosaic distributions of six families of transposon-derived transcription factors (TFs) across metazoans. All TFs depicted in the tree are lineage-specific genes that have no homolog in other classes or phyla. Each colored circle represents one of the six TF protein families: ZBED, CNPB, FHY3, HTH-Psq, THAP, and FLYWCH. Figures within circles indicate several independent species-specific events of the domestication of a particular TF family. The total numbers of transposon-derived TFs identified in each reference species are shown on the right. We observed the most extensive expansion of transposon-derived TFs in four bilaterian lineages led to the hydrozoan polyp - *Hydra* (142), the oligochaete - *Capitella* (98), the sea slug - *Aplysia* (59), and the bivalve - *Crassostrea* (91). Of note, a significant expansion of the THAP gene family occurred in *Capitella* (87), *Hydra* (73), and *Crassostrea* (58). Independent species-specific expansions of the FLYWCH gene family occurred in ctenophores *Mnemiopsis* (16) and *Pleurobrachia* (16).

**Figure 2:**
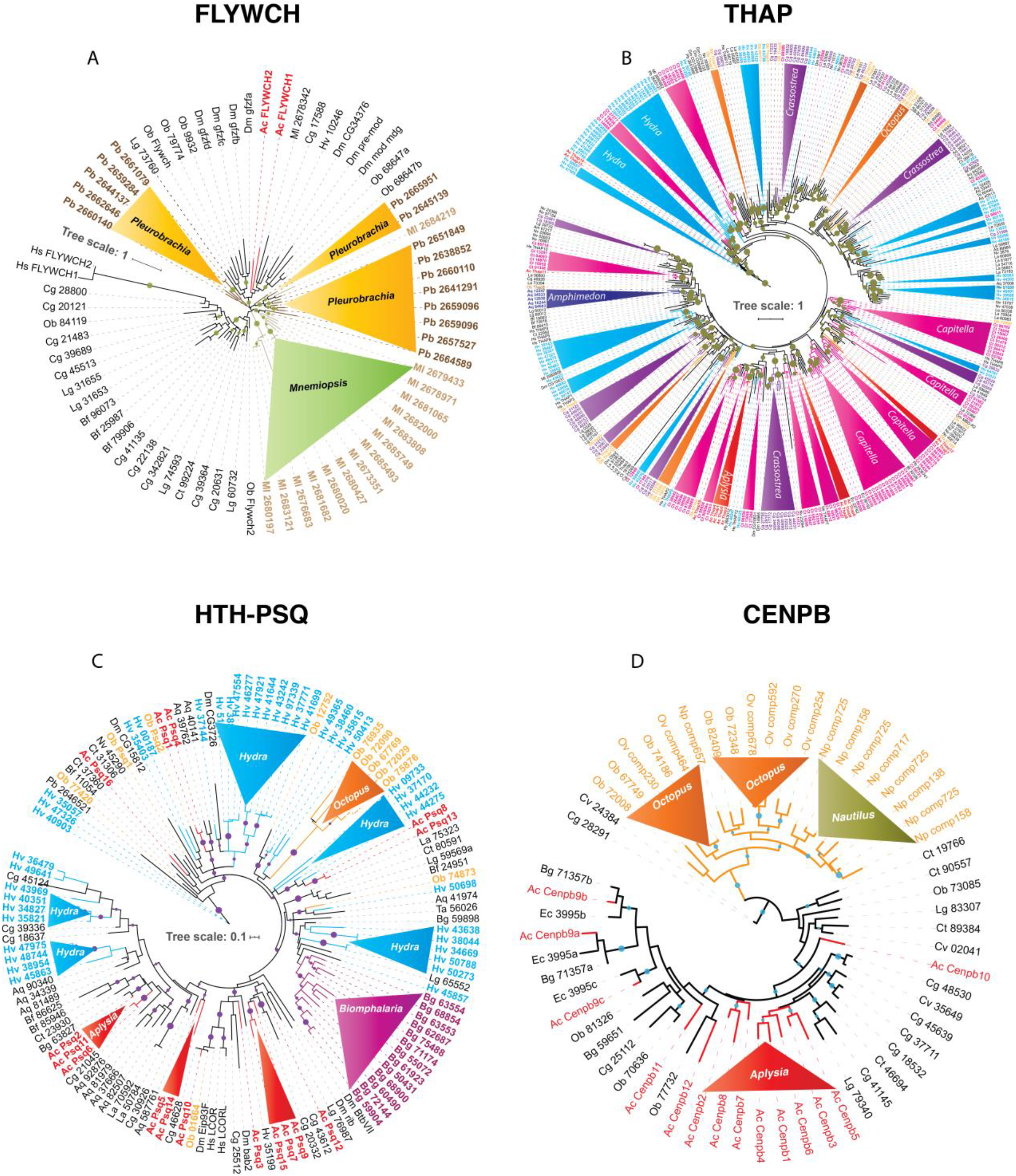
Independent expansion and convergent evolution of transposon-derived transcription factors in Metazoa. The phylogenetic tree represents the independent expansion and evolution of transposon-derived transcription factors protein families across metazoans. Each solid-color triangle represents species-specific expansion that has no homologs in related species. We used the following DNA binding domains–FLYWCH (A), THAP (B), HTH-PSQ (C), and CENPB (D) - as illustrative examples to build the maximum likelihood (ML) tree. The trees show independent FLYWCH gene expansion in the ctenophores *Mnemiopsis* and *Pleurobrachia* (A). Similarly, independent THAP genes expansion in *Capitella, Octopus, Crassostrea, Hydra* (B), HTH-Psq expansion in *Hydra, Biomphalaria, Aplysia*, and *Octopus* (C), and Independent convergent domestication of CENPB genes in *Octopus, Nautilus*, and *Aplysia* (A). High-resolution images of each of these trees are presented in Figs. 1S-4S.

**Figure 3:**
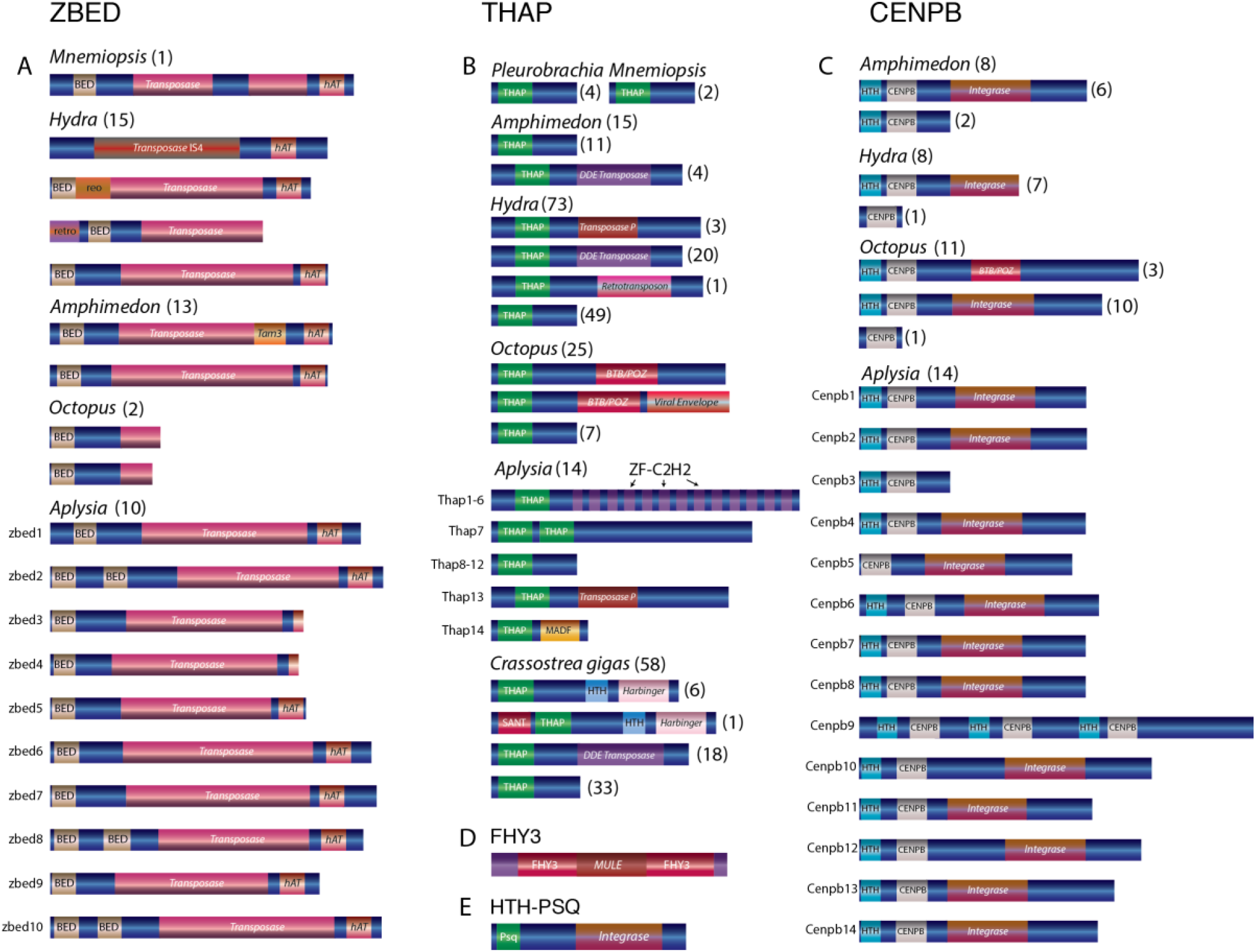
Domain organizations of the transposon-derived transcription factors across metazoans. Transposon insertions domains are shown in shaded red color and labeled as *integrase, transposase, Harbinger, BTB/POZ*, etc. Note that the same transcription factor protein families have different transposon components. For example, *Octopus* CENPB and THAP proteins have derived mostly from BTB/Poxvirus (BTB ^78^/POZ^79^) transposable elements, whereas, in other species, the same TFs have originated from multiple different transposable elements. Similarly, *Hydra* ZBED genes could have derived from at least three transposon sources such as *retrotransposon, reoviruses*, and *transposon IS4*, whereas all *Aplysia* ZBED genes seem to have derived from *Ac* transposon (**Fig. 5S** and **6S**). Numbers within the parentheses indicate the number of genes for each particular domain organization.

All TE-derived TF families identified in our analysis showed low (<1; Z-test *P*<0.05) nonsynonymous substitutions versus synonymous substitution (Ka/Ks) ratios, indicating negative or purifying selection acting to maintain evolutionarily conserved sets of amino acid sequences. Similarly, the low Ka/Ks ratio of TE-derived TFs also suggests stationary domesticated genes ^41^. In addition to the Z test, Fast Unbiased Bayesian Approximation (FUBAR) ^42^ estimation of the dN/dS ratio also confirmed negative or purifying selection pressure acting on these TFs (**Fig. 4**). The total number of the proposed transposon-derived TFs is 788. We include species such as the sea slug, *Elysia chlorotica*, the hemipteran insect *Myzus persicae*, and the rainbow trout *Oncorhynchus mykiss*.

**Figure 4:**
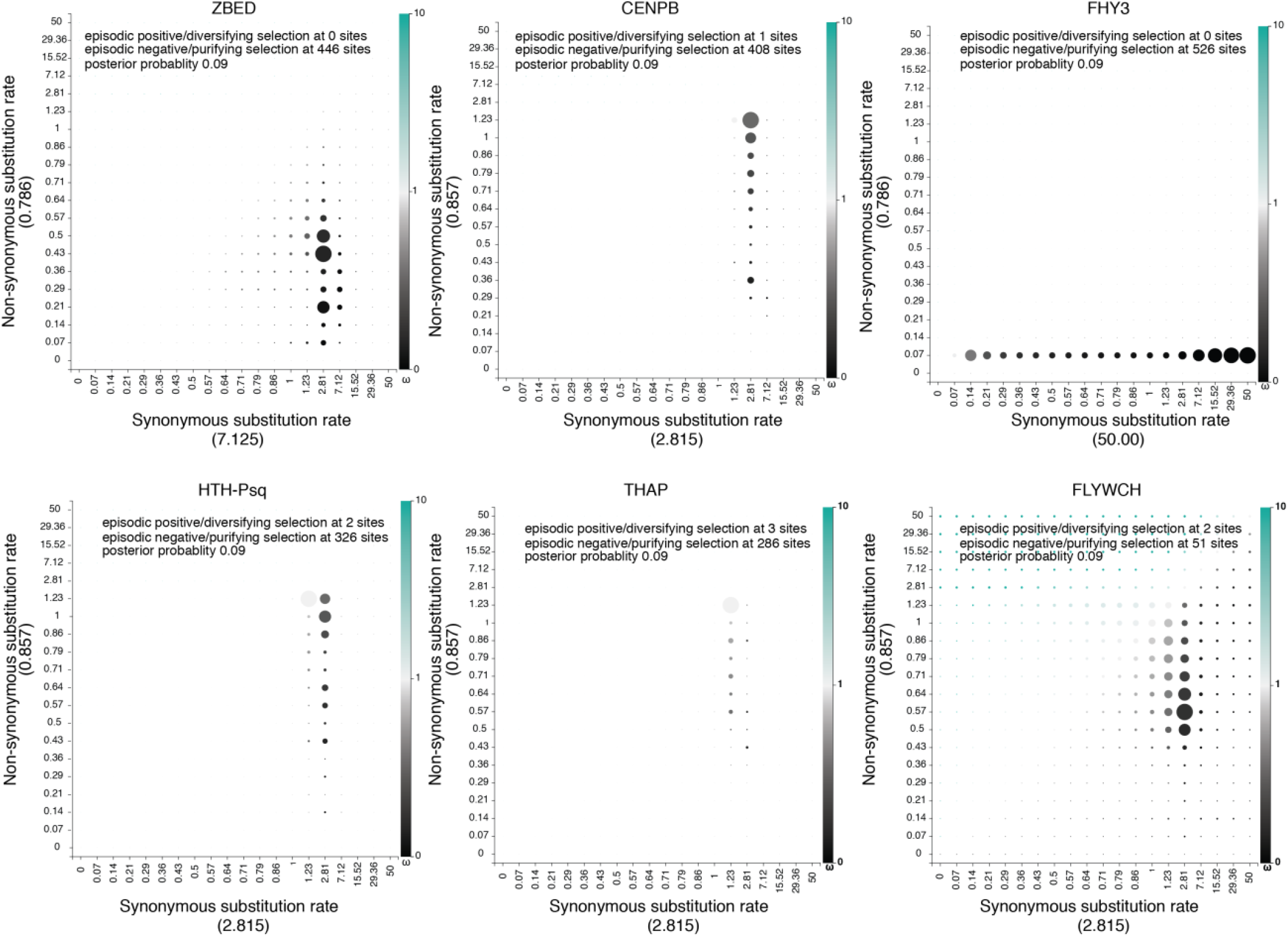
Non-synonymous (dN) versus synonymous substitution (dS) ratio show transposon-derived transcription factors evolving under purifying selection pressure. Non-synonymous versus synonymous substitutions were calculated across all TE-derived TF families using the Fast Unbiased Bayesian Approximation (FUBAR) approach ^42^. Synonymous substitutions (dS) rates calculated under each family showed in X-axis inside the parentheses. Similarly, Non-synonymous substitutions (dN) rates calculated under each family showed in Y-axis inside the parentheses. Gray to intense black color-coding dots signifies negative or purifying (dN/dS <1) selection, while light green to intense green represents sites under diversifying or positive (dN/dS >1) selection.

**Fig. 1** illuminates the mosaic-type distribution in the recruitments of transposon-derived TF subfamilies across major metazoan lineages studied here. In the sister group to all Metazoa – Choanoflagellata – we found only two genes likely encoding transposon-derived TFs from ZBED and THAP superfamilies, respectively.

Ctenophores are viewed as the earliest branching lineage of animals, sister to the rest of Metazoa ^43-46^, although the reconstruction of the basal metazoan phylogeny is still a highly debated topic ^47-50^. Unlike other studied metazoans, both *Mnemiopsis* and *Pleurobrachia* showed tremendous expansions of the FLYWCH transcription factor gene family (**Fig. 2A**). FLYWCH ^51,52^ is a distinct DNA-binding zinc finger domain-containing protein family known to have originated from the *Mutator* transposase ^53^. FLYWCH domains are evolutionary conserved but relatively rarely occur in animals. They were initially identified in *Drosophila* ^54^ and then in *C. elegans*, where it plays regulatory roles during embryogenesis by repressing microRNAs ^52^. The most recent evidence suggests that **FLYWCH**, in complex with β-catenin, repressed particular genes of the Wnt pathways and, therefore, can control cell polarity, migration, and metastasis ^55^. Surprisingly, none of the newly identified ctenophore FLYWCH domain-containing genes have homologs in each other ctenophore species (**Fig. 2A**). Unfortunately, there are no functional studies of these genes, and the roles of these TFs in ctenophores would be subjects of future studies.

There are three species with the broadest overall domestication of TEs: the hydroid polyp – *Hydra* (149 TFs), the polychaete annelid – *Capitella* (102 TFs), and the gastropod mollusk, *Aplysia* (100 TFs). In these animals, the identified domestication events are both species-specific and TF-type-specific. In other words, for each animal studied, we noticed an independent expansion of one or more families of TE-derived TFs (**Fig. 1**). The most notable examples of TE exaptation we found in *Hydra* and the ctenophore *Pleurobrachia* (5 out of 6 superfamilies), *Aplysia* (6 out of 6 superfamilies), and the sponge *Amphimedon* (5 out of 6 superfamilies). Surprisingly, the lineage that led to the sponge also revealed multiple examples of independent domestication and expansion of TE-derived TFs compared to other non-bilaterian metazoans (except *Hydra*), which correlate to astonishing diversification within the phylum Porifera in general.

In contrast, the placozoan *Trichoplax* – the simplest known free-living animal ^56,57^, had the smallest number (9) of TE-derived TFs, which might reflect the observed morphological simplicity of these disk-shaped benthic animals with only three layers of cells gliding on algal substrates ^57-60^.

Likewise, the anthozoan *Nematostella* also had a modest representation of TE-derived TFs (21), mostly related to just one superfamily; there are 15 **Th**anatos and **a**ssociated **p**rotein (**THAP**) domain-containing genes. THAP genes were found in *Drosophila*, and they are known to have originated from **P** element transposes ^61^. Our analysis support events of the independent diversification of THAP genes in *Hydra* (73), *Capitella* (87), *Crassostrea* (58) (see details in the next section and **Fig. 2B**; Fig. 2S); and at a lesser degree in a living fossil - the brachiopod, *Lingula* (27) and *Octopus* (25).

In summary, THAP genes represent the largest class of TE-derived TFs identified in this study, including the basally branched chordate amphioxus (*Branchiostoma*) and humans. THAP-TF functions in invertebrates are primarily unknown ^62^. On the other hand, THAP TFs in humans were implicated in epigenetic regulation, maintenance of pluripotency, transposition, cancers, and other disorders like hemophilia. For example, THAP0 is a member of the apoptotic cascade induced by IFN-γ ^63^. THAP1, with RRM1, regulates cell proliferation ^32^. THAP5 acts as a cell cycle inhibitor ^33^. THAP9 is an active transposase in humans ^31^. The THAP11 homolog in mice is essential for embryogenesis in mice ^64^.

Two other groups of presently identified TE-derived TFs are also prominent in humans and *Branchiostoma*: ZBED and CENPB (**Fig. 1**).

**BED z**inc fingers or **ZBED** genes reported having derived from the *hAT* (***h****obo*, ***A****c*, ***T****am3*) superfamily of DNA transposon ^65^, and members of this superfamily regulate an extensive array of functions in vertebrates. For example, ZBED6 affects development, cell proliferation, wound healing, and muscle growth ^30^. ZBEDs are present in mammals, birds, reptiles, and fish; however, they are absent from jawless fishes. Based on these findings, it was proposed that ZBED genes in vertebrates originated due to at least two independent *hAT* DNA transposon domestication events in primitive jawed-vertebrate ancestors ^29^. Our searches against the *Branchiostoma belcheri* genome uncovered a full-length ZBED gene, which was surprisingly absent from the *Branchiostoma floridae* genome, further indicating species-specific and mosaic exaptation of TE-encoded genes.

Also, using both the DNA binding BED domain and known full-length ZBED genes, we find that ZBED genes form a monophyletic cluster in three mollusks (*Aplysia, Biomphalaria, Crassostrea*), the sponge *Amphimedon*, and *Hydra* (**Fig. 5-6S**). **Cen**tromere-binding **p**roteins-**B** (**CENPB**) transcription factor ^66^ involved in chromosome segregation maintenance and genome stability ^67^ recurrently domesticated from *pogo*-like transposons ^14,15^ across Metazoa (**Fig. 7S**). CENPB homologs were found in mammals ^68^ but not in other vertebrates. Nevertheless, we identified CENPB TFs from both *Branchiostoma belcheri* and *B. floridae* genomes, indicating their presence before the divergence of vertebrates. Thus, this finding suggests either loss of CENPBs in most of the extant lineages of vertebrates or their independent domestication in mammalian species, which is a more likely scenario ^14^. There is also a remarkable diversification and independent expansion of the CENPB superfamily in Mollusca (**Fig. 7S**), which we will discuss in the following section.

The most stunning example of mosaic recruitment of TEs can be illustrated using *Mule* transposons. *Mule* transposon-derived transcription factor **f**ar-red elongated **hy**pocotyls 3 **(FHY3)** group are critical for far-red (near-infrared) light signaling and survival of chloroplast in plants ^26,69^. Here for the first time, we identified FHY3 in animals (**Figs. 1, 3D**). Our cross-species comparison across metazoans showed that FHY3 was present in three copies, both in the demosponge *Amphimedon* and the sea slug *Aplysia* genomes. There are two copies in the brachiopod *Lingula* and one in *Octopus* genomes (**Fig. 1**). However, we did not find FHY3 in the sequenced ctenophore (*Pleurobrachia* and *Mnemiopsis*), placozoan (*Trichoplax*), and cnidarian (*Nematostella* and *Hydra*) and human genomes. Thus, FHY3 can be absent or present in a mosaic fashion without a recognized taxonomical specification. Our phylogenetic analysis (Excel File 1S) showed that FHY3 had been repeatedly domesticated over 550+ million years of animal evolution (see **Fig. 8S**), including examples from selected mollusks (e.g., the algae-eating sea slugs *Aplysia californica, Elysia chlorotica*, and the oyster - *Crassostrea*), some arthropods (*Myzus persicae* and *Limulus polyphemus*) and chordates (*Branchiostoma*).

In conclusion, we obtained robust evidence that the majority of TFs are the results of the species-specific convergent domestication events across all animal phyla tested. **Fig. 2** illustrate these cases. Of note, although some of the studied species show a predominant exaptation of just one or two categories of genes, many domesticated events occurred independently, even within the same superfamily of TE-derived TFs (**Figs. 2, 1S-8S**). This situation is summarized below, focusing on the Lophotrochozoan lineage.

### 2. Transposon-derived TFs showed independent species-specific expansion and evolution in Molluscs

Lophotrochozoa or Spiralia, including the phylum Mollusca, is the most morphologically and biochemically diverse animal clade ^36,70,71^. None of the identified TE-derived TFs were previously reported in Lophotrochozoa (Table 1). The phylum Mollusca in our analysis is represented by seven species (*Aplysia, Biomphalaria, Elysia, Lottia, Crassostrea, Octopus*, and *Nautilus*), with *Aplysia* showing the most remarkable expansion of TE-derived TFs (**Fig. 1**).

First, we systematically scanned the complete set of the TFs encoded in the *Aplysia californica* genome (a prominent neuroscience model^72^), resulting in the identification of 824 transcription factors. Then, we identified 59 novel (∼7%) transposon-derived TFs that have no homolog in closely related species such as in *Biomphalaria* (the freshwater pulmonated snail ^73^) or the limpet *Lottia* ^74^. This finding indicates that these TFs were not originated from canonical gene duplication events (Excel File 1S); they do not follow the canonical subfunctionalization ^75^ and neofunctionalization ^76^ characteristics. Of these 59 *Aplysia* lineage-specific TFs, 42 were coupled with the transposase (TPase) domain (**Fig. 3**), confirming the hypothesis that these genes, including their DNA-binding domain, may have originated by unique mechanisms involving ‘*cut-and-paste’* DNA transposons.

In mollusks, we also revealed that the lineage-specific TFs, even those belonging to identical TF families, originated both from similar and different transposon sources: the majority of TE-derived TF domestication events were not detected from related species. Thus, the most likely parsimonious scenario is a broad scope of independent domestication events leading to convergent evolution of TE-derived TFs within animal lineages studied here. **Fig. 2** illustrates bursts of parallel expansions of transposon-derived TFs subfamilies. Three examples are outlined below.

There are convergent domestications of *pogo*-derived CENPB sequences in *Aplysia*, cephalopods, and other Lophotrochozoan species such as in *Crassostrea* (**Fig. 2D**). Within the cephalopod lineage, we identified two distinct events of *pogo* domestication—one, in the lineage leading to *Nautilus* and another event occurring in the lineage leading to *Octopus* (**Fig. 2D**).

**H**elix-**t**urn-**h**elix motif of **p**ip**sq**ueak (**HTH-Psq**) proteins form a family of transcription factors known to have derived from *Drosophila* pogo transposase ^77^. We find the *Aplysia* genome encodes 16 **HTH-Psq** subfamily transcription factors while the *Biomphalaria* genome encodes 15 of them. Surprisingly none of these *Biomphalaria* TFs has direct homologs in the *Aplysia* genome and vice versa (**Fig. 2C; Fig. 3S**), indicating species-specific expansion event. Similarly, both *Hydra* and *Octopus* showed independent species-specific expansions of transposon-derived HTH-Psq genes. Thus, independent domestication of Psq genes might occur at least five times in the genomes of *Aplysia, Biomphalaria, Octopus*, and the *Hydra* and *Amphimedon* genomes (**Fig. 2C**).

**M**yb-SANT, like in **Adf** (**MADF**) domain-containing genes initially identified in *Drosophila* known to have originated from the **P i**nstability **f**actor or PIF superfamily of DNA transposon ^26^. We find that MADF genes were expanded in *Amphimedon, Drosophila*, and, most of all, *Aplysia* with at least six predicted independent domestication events (**Fig. 4S**). Although MADF genes likely derived from PIF superfamily of DNA transposon, we have excluded MADF genes owing to growing concern that these genes do not harbor a recognized transposon-derived conserved domain within the protein-coding gene.

Altogether our results suggest that there was a substantial lineage-specific diversification, and independent evolution of new genes originated from a modular diversity of *cut-and-paste* DNA transposons, as we outlined in the next section.

### 3. Domain analysis revealed the presence of transposons derived components within the protein-coding TFs

All subfamilies of transposon-derived TFs predicted in this analysis have a modular domain architecture (**Fig. 3)**. Within each subfamily most TFs encoded recognizable transposon-derived components within exons of these protein-coding genes. For example, transposon-derived ZBED TFs, besides encoding the canonical DNA-binding BED zinc finger motif, also encoded a transposon-derived *transposase* domain and a *hAT* dimerization domain (**Fig. 3A**). Strikingly, we find that ZBED genes across metazoans derived from diverse transposable element components. For instance, *Homo* ZBED5 is known to have derived from *Buster* DNA transposon ^29^, which in our analysis forms a robust clade with one of the *Octopus* ZBED genes indicating its *Buster* transposon origin (**Fig. 5S & 6S**). In contrast, the second *Octopus* ZBED gene forms a robust cluster with the *Hydra retrotransposon*-derived ZBED gene (**Fig. 5S & 6S**). This result indicates that the two *Octopus* ZBED genes may have evolved from two independent transposon components.

Similarly, *Hydra* ZBED genes contained at least three transposon components, such as *retrotransposons, reoviruses*, and *transposon* IS4 (**Figs. 3A, 5-6S**). Likewise, while *Octopus* THAP genes are mostly derived from BTB ^78^ (Broad-Complex, Tramtrack, Bric a Brac) or POZ ^79^ (poxvirus and zinc finger) transposon sources — the *Hydra* THAP genes, however, found to be derived from versatile transposon sources such as *Transposase P* element, *DDE transposase* (*DDE_Tnp_4*) and *retrotransposon*. In contrast, some of the *Crassostrea gigas* THAP genes contained sequences associated with the *Harbinger*-derived transposon domain (**Fig. 3B**). Also, while most of the *Octopus* CENPB TFs showed an association with the transposon-derived BTB/POZ domain, none of the genes from another mollusk, *Aplysia*, contained this domain (**Fig. 3C**).

Both CENPB and HTH-Psq genes had a signature of the viral *rve* superfamily of the retroviral integrase domain (**Fig. 3C, E**). Integrase is the retroviral enzyme that catalyzes the integration of virally-derived DNA into the host cell’s nuclear DNA, forming a provirus that can be activated to produce viral proteins ^80^. In the same way, FHY3 genes share remarkable sequence similarities with MURA ^81^, the transposable element encoded by the *Mutator* element of maize, and the predicted transposase of the maize mobile element *Jittery* ^82^. Both of these transposons are a member of the **Mu**tator-**l**ike **e**lements (*MULE*) ^83^(**Fig. 3D**).

These results, for the first time, indicate that even within the same subfamily of transposon-derived TFs — similar domains have derived from multiple transposon components across the animal kingdom. Together our phylogenetic analysis and the revealed domain organizations suggest that similar domain architecture originated in parallel from numerous transposon resources across phyla.

## SUMMARY

By systematic analysis of seven thousand animal TFs, we have identified a total of 788 (>10%) novel DNA transposons-derived TFs across metazoans (**Fig. 1**; Excel File 1S). Overall, these TFs show mosaic patterns in their distribution with extreme heterogeneity and with a ‘sudden’ appearance in one lineage and, at the same time, found to be ‘missing’ in more closely related species. Although the majority of studied species predict a predominant exaptation of just one category of genes, many domesticated events might occur independently in evolution, even within the same superfamily of TE-derived TFs (**Fig. 2**). These results suggest a substantial lineage-specific diversification, and independent origins of new TF genes originated from a broad array and a modular diversity of *cut-and-paste* DNA transposons, and perhaps related viroid-like elements. Many of the described here TFs preserved the original modular gene organization (**Fig. 3**) and could act as highly dynamic modules shaping the genome-wide reorganization within Metazoa.

### Methods

All methods are summarized in the Method section at next pages

## Methods

### Genome-wide identification and annotation of transcription factors across metazoans

We used experimentally verified DNA binding domains as a probe (query) to retrieve the complete repertoire of the transcription factors encoded in the metazoan genome. For example, using the entire dataset of experimentally verified 1600 transcription factors encoded in the human genome as a query^38^, we manually curated, annotated, and identify that the *Aplysia* genome encodes 824 transcription factors. Similarly, the full dataset of transcription factors from both sea slug *Aplysia* and humans was used to retrieve a complete set of transcription factors from *Octopus bimaculoides* and other metazoans such as sponge *Amphimedon, Trichoplax*, and ctenophore *Pleurobrachia*, etc. (data not shown). While annotating the *Aplysia* genome for the transcription factors we noticed that the *Aplysia* genome encodes a substantial number of the transposon (TE) derived transcription factors (TFs) (100/824 = ∼13%), which prompted us to screen the presence of TE-derived TFs across metazoans.

### Identification of transposable element-derived transcription factors

We performed comprehensive searches for the transposable element (TEs) - derived transcription factors (TFs) in representatives of all five metazoan lineages such as Ctenophora, Porifera, Placozoa, Cnidaria, and Bilateria, including its three superclades (Deuterostomia, Escdysozoa, and Lophotrochozoa)^36,43,44,84^. The Method section summarizes the species used in this study and their reference genomes.

**Table 1S.**
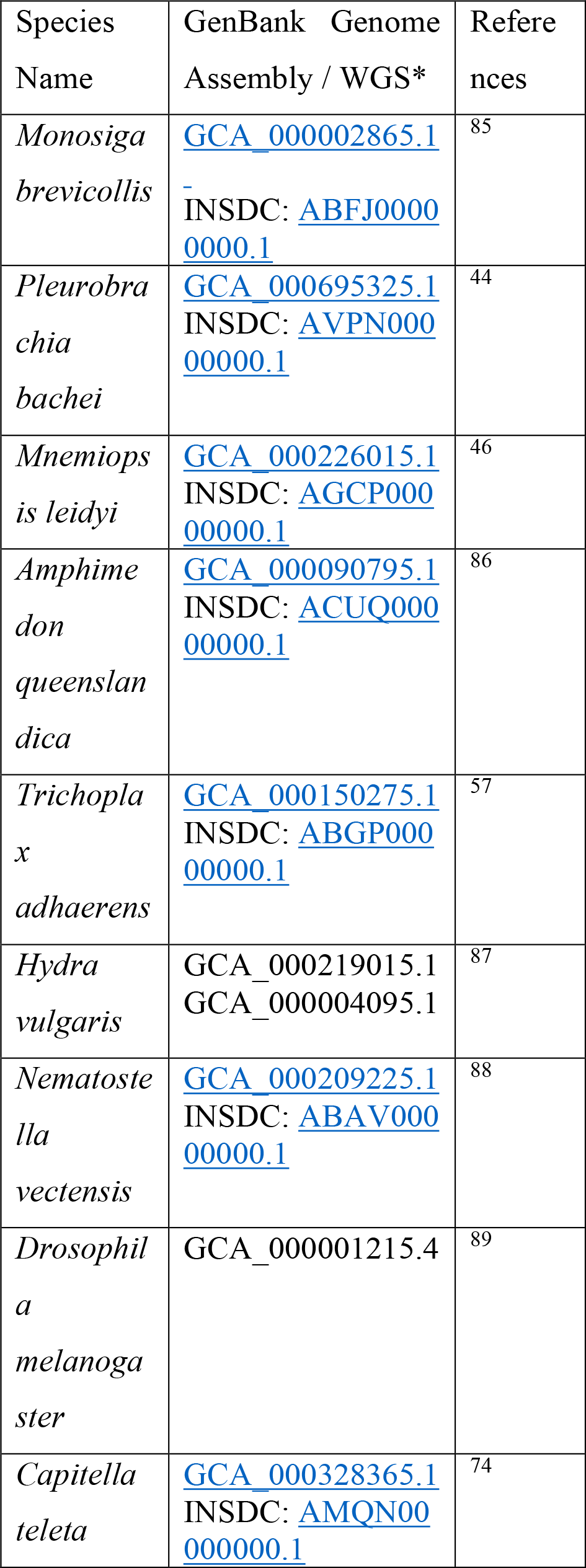
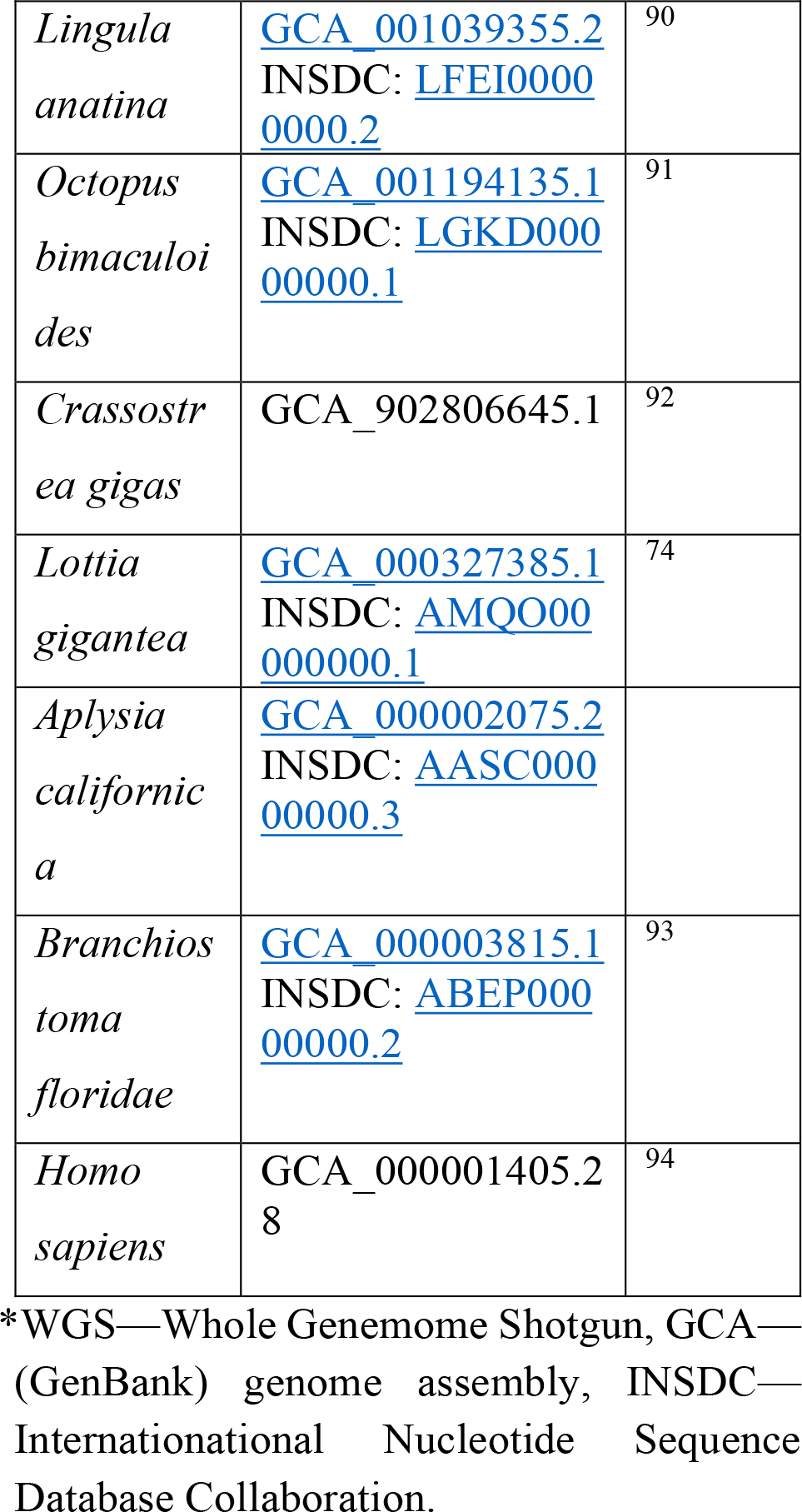
Species used in comparative analyses.

We used representatives of published and confirmed domesticated transposable element-derived TFs protein families from both plants and animals as a query. Both PSI-BLAST, as well as Tblastn searches, were performed using both the command-line version at the NCBI standalone BLAST (version 2.2.18)^95^ as well at the on-line BLAST web interface^96 97^using default e-value cut off for the online version and 10^−5^ to 10^−10^ cut off for the stand-alone blast to identify all potential homologs. Homologs were detected not solely based on e-value cut off but other criteria such as coverage statistics, bit score, etc. were considered. Protein sequences recovered from one round of TBLASTN or PSI-BLAST searches were recursively used as queries until no further sequences were detected. Each protein blast hit was manually inspected following multiple sequence alignment (MSA) and validated utilizing several databases including the NCBI conserved domain database (CDD)^98^, Hmmer^99^, Pfam^100^ and SMART^101^. In the case of non-availability of the gene model (exome), genome sequences surrounding the coding region were excised, and homology-based gene prediction based on hidden Markov models (HMMs) was performed in FGENESH+ (www.softberry.com) to identify the complete open reading frame. Finally, TE insertions within the TFs were further validated by similarity searches against the *de-novo* assembled RNA-Seq (transcriptome) datasets obtained in Moroz lab (https://neurobase.rc.ufl.edu).

**Table 2S:**
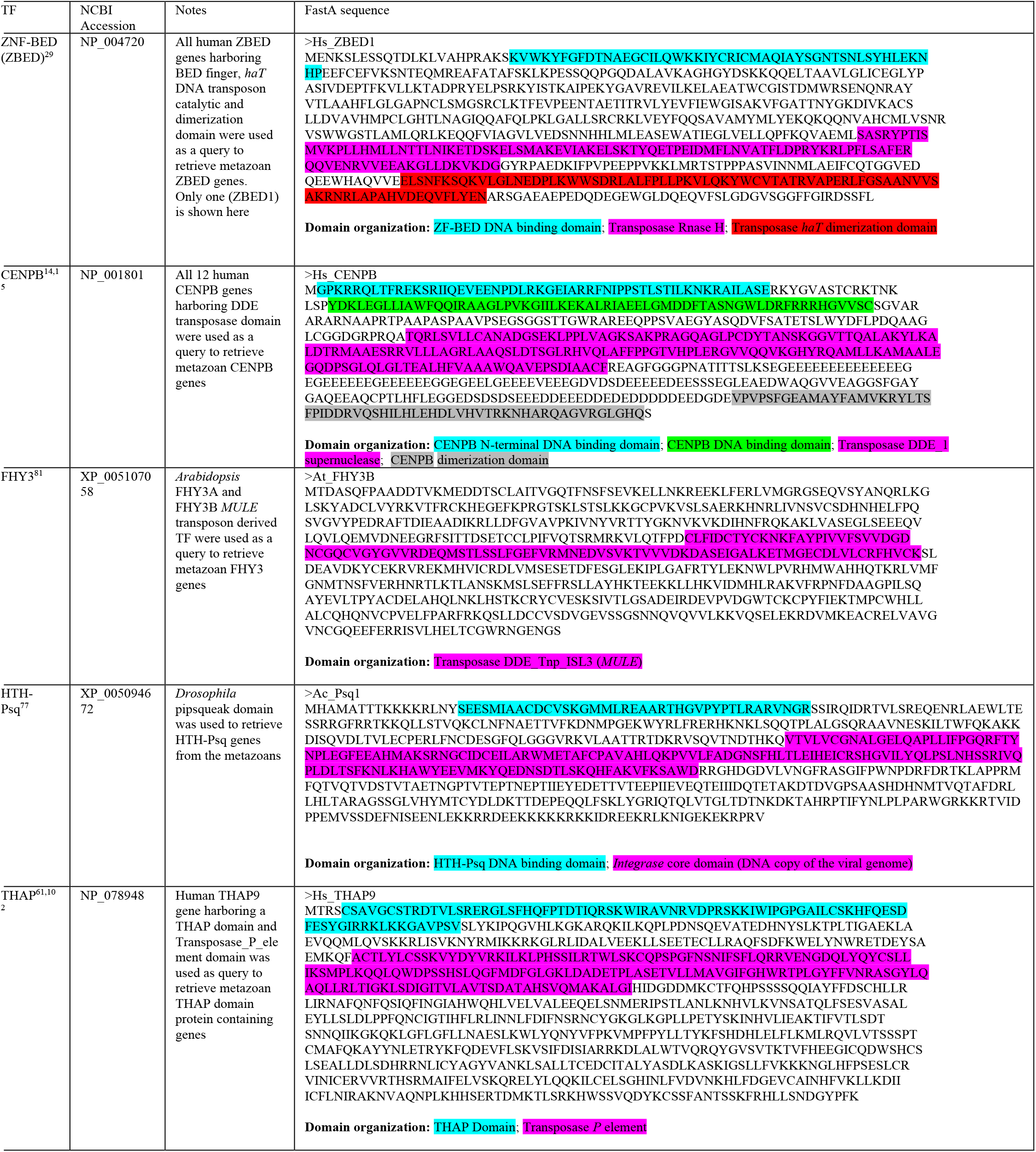
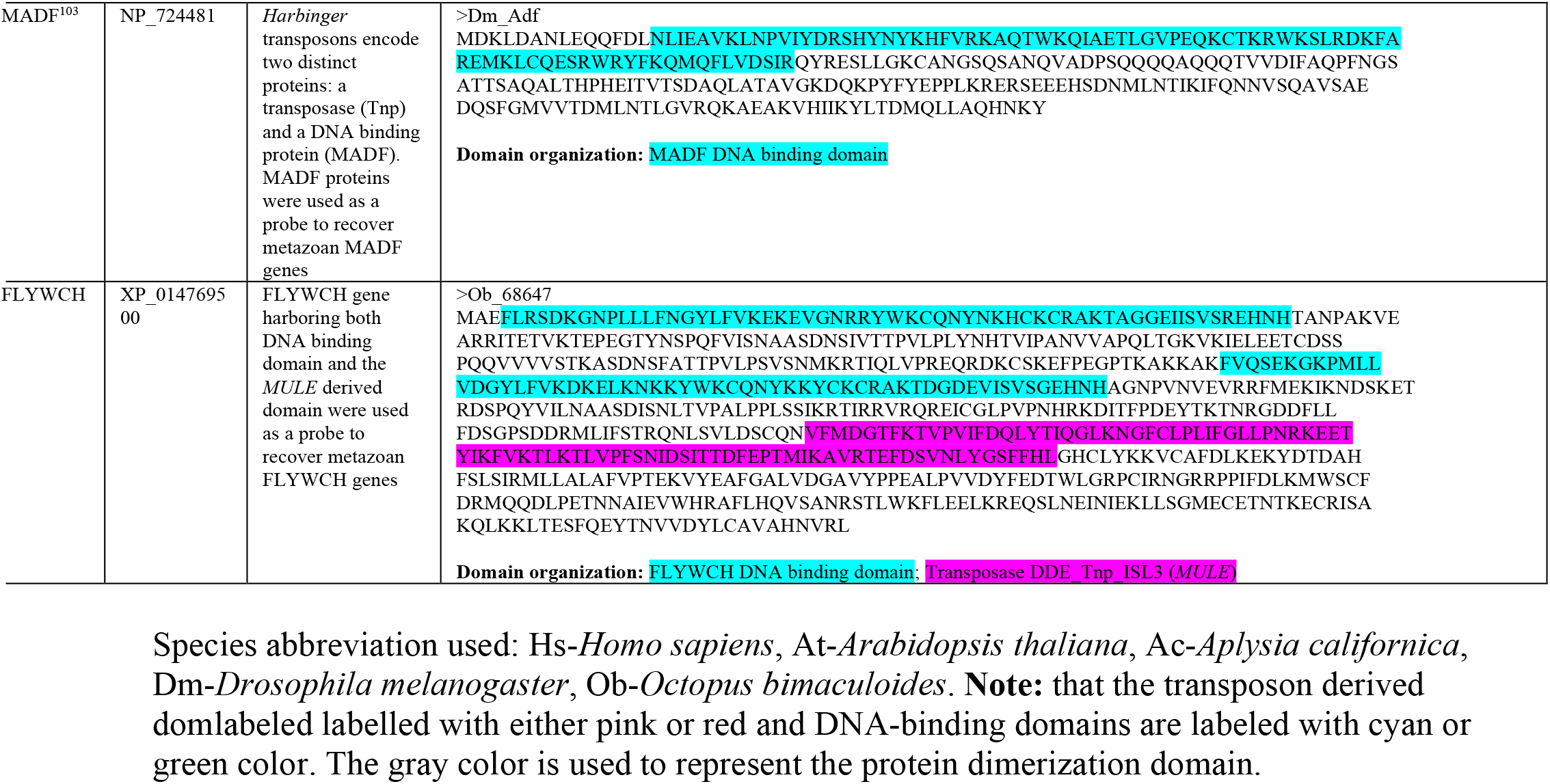
Experimentally verified and published transposon-derived transcription factor proteins were used as a query to search against the metazoan genomes.

### Multiple alignment and protein domain identification

Protein functional domains were identified by sequence search of the NCBI conserved domain databases^98,104^. Results were verified via sequence searches of the SMART^101^ and Pfam database^105^. Also, sequences were aligned in MUSCLE^106,107^ and displayed in clustalX^108^ and manually confirmed the domain architecture by examining the sequences based on protein secondary structure analysis and profile alignments. Multiple sequence alignment (MSA) obtained through MUSCLE was used to build the HMMER v3.1b2^99^ position-specific scoring matrix (PSM) to search against the reference proteome datasets.

### Phylogeny

Maximum-likelihood (ML) trees were inferred using PhyML v3.0^109,110^, with the best-fit evolutionary model identified using the AIC criterion estimated by ProtTest^111^. ML phylogenies were performed using the JTT model of rate heterogeneity, estimated proportion of invariable sites, four rate categories, and estimated alpha distribution parameter. Tree topology searches were optimized using the best of both NNI (nearest-neighbor interchanges) and SPR (subtree pruning and regrafting) moves^112^. Clade support was calculated using the SH-like approximate likelihood ratio test^113^. Unless otherwise mentioned, all phylogenetic trees presented throughout the manuscript showing SH-support of 80 or greater. The resulting phylogenetic trees were viewed and edited with iTol version 2.0^114^.

### Codon substitution pattern and inference of selective pressures

Protein sequences of TE-derived transcription factors under each family were aligned using MUSCLE^107^ and the conversion of protein alignments to corresponding nucleotide coding sequences were obtained using PAL2NAL webserver^115^. Codon-based tests of neutrality and negative or purifying selection were conducted using MEGA with a Z test by calculating the substitution ratio of the number of non-synonymous substitution per non-synonymous site (Ka) versus synonymous substitution per synonymous sites (Ks) using the Nei-Gojobori method^116^. Orthologous sequences with a Ka/Ks value of <1 (Z-test, *P* <0.05) were defined as having been under purifying selection shown with yellow color.

Alternatively, we used the Bayesian approach of Fast Unbiased Bayesian Approximation (FUBAR)^42^ datamonkey.org) to infer nonsynonymous (dN) and synonymous (dS) substitution rates on a per-site basis on a given codon alignment and corresponding phylogeny (**Fig. 4**).

## Data Availability

The data that support the findings of this study are available from the corresponding author upon reasonable request.

## Author contributions

K.M. and L.L.M.: Conceptualization; Writing an original draft, Writing-review & editing, Data obtaining, and curation. K.M: Formal computational analysis, Investigation, Methodology, Software, Validation, Data Visualization, L.L.M.: Funding Acquisition, Project Administration, Resources, and Supervision.

## Conflict of Interests

The authors have no conflict of interest to declare.

## Acknowledgments

The authors would like to thank Drs. Caleb Bostwick, Peter Williams, and Andrea Kohn for the generation of RNA-seq libraries and initial annotations. Thanks to Gayle Prevatt for the initial drawing of the animal sketches. This work was supported by the Human Frontiers Science Program (RGP0060/2017) and National Science Foundation (grants 1146575, 1557923, 1548121, and 1645219 to L.L.M). The research reported in this publication was also supported in part by the National Institute of Neurological Disorders and Stroke of the National Institutes of Health under Award Number R01NS114491 (to L.L.M.).

## Notes

### Competing Interest Statement

The authors have declared no competing interest.

